# A Perfect Soldier: the black soldier fly as a microbial-mediated physiological resilience model

**DOI:** 10.64898/2026.01.18.700162

**Authors:** Hunter K. Walt, Michael B. Smith, Erin Harris, Sienna McPeek, Florencia Meyer, Spencer T. Behmer, Federico G. Hoffmann, Jeffery K. Tomberlin, Christine J. Picard, Heather R. Jordan

## Abstract

Understanding the complex interplay between a host, its diet, and its microbiome is crucial for comprehending an organism’s health and adaptability. Diet impacts both the host and microbiome, which then influence each other. We used black soldier fly larvae (*Hermetia illucens*) as a model to investigate this tripartite interaction due to its resilience and bioconversion capabilities. We analyzed life-history traits and metatranscriptomics in larvae fed three diets: carbohydrate-rich, protein-rich, and balanced. Our results showed that dietary macronutrients correlated with shifts in the microbial community and gene expression. The carbohydrate-rich diet, in particular, led to increased microbial diversity and carbohydrate metabolism transcripts. However, this diet also negatively affected larval weight and development, suggesting potential host control over the microbiome. Overall, black soldier fly performance was highest on the balanced diet. This study highlights the black soldier fly’s resilience and its value as a model for exploring host-diet-microbe interactions.

**Significance Statement:** Understanding the intricate interplay between an organism, its diet, and its microbiome is fundamental to health and adaptability. This complex tripartite relationship, where dietary macronutrients influence microbial communities and their gene expression, while the host maintains control, is crucial for addressing global challenges from sustainable food systems to personalized medicine. Using the black soldier fly as a resilience model, our metatranscriptomic study reveals how specific dietary shifts impact both host and microbial gene expression, providing mechanistic insights into nutrient utilization and adaptability with broad implications for diverse biological systems, including animals and humans.

## Introduction

Few interactions are as fundamental as the interplay between an organism, its diet, and the vast community of microbes living within and on it. This tripartite interaction is extremely important, shaping everything from how we digest our food and bolster our immune systems to our overall health and fitness. However, the host-diet-microbe relationship is complex, as it is characterized by the interplay between microbial competition over these dietary resources while the host keeps its microbiome under control. Unraveling this complex interplay could be key to addressing some of humanity’s most pressing challenges, from creating sustainable food systems to developing personalized medicine (1).

Understanding how specific dietary macronutrient biases, particularly in the case of the protein-to-carbohydrate ratio, shape the gut microbial community structure and their gene expression is of paramount importance (2). This research is crucial because the gut microbiome is a key determinant of host health and fitness, mediating a vast array of physiological processes, from nutrient absorption and immune system development to protection against pathogens (3).

Understanding how diet influences not just the diversity and abundances of microbes present, but also their functional output (i.e., which genes they are expressing and what metabolites they are producing), provides a mechanistic link between diet and host outcomes. This complex interplay extends beyond simple digestion, fundamentally influencing host metabolism, immunity, and even neurobehavioral pathways, making it a central pillar of precision nutrition and personalized medicine (1).

At its core, understanding this host-microbe-diet axis means grasping not just which microbes are present, but also what they are doing and how their actions affect the host. While we have long known that diet influences the gut microbiome (2, 3) the cutting edge of research lies in tracing the intricate molecular conversations that unfold. How does a diet rich in protein or carbohydrates alter the gene expression of specific microbial groups? And how do these microbial changes, in turn, influence the host’s own gene expression, metabolism, and even its susceptibility to disease? To decode these dynamics, researchers are moving beyond simple snapshots of microbial diversity and employing more advanced techniques like metatranscriptomics. This powerful technique allows us to simultaneously identify the microbes present, quantify their abundance, and reveal which of their genes are actively being expressed (4). Additionally, metatranscriptomic sequencing captures the host’s gene expression at the same time, offering a holistic view of how both the host and its microbiome respond to dietary shifts. This comprehensive, systems-level approach is crucial for pinpointing the causal links and predictive markers that define diet-driven host-microbe interactions.

To unravel complex relationships, scientists often turn to model organisms that offer unique insights. To study diet-host-microbe tripartite interactions, the black soldier fly (*Hermetia illucens*) stands out as an exceptionally compelling subject. This insect is a remarkable generalist, capable of thriving on an astonishingly diverse array of organic matter, from decaying plants to food waste (5). This extraordinary dietary flexibility makes it an ideal system for studying how organisms adapt to wildly fluctuating nutrient availability. Beyond its biological intrigue, the black soldier fly is also gaining significant industrial importance (6). It is a champion of bioconversion, efficiently transforming organic waste into high-quality protein and fats, offering a sustainable source of nutrients for animal feed and even human food (7).

The black soldier fly’s impressive resilience, a hallmark of generalist species, is deeply intertwined with its broad dietary range (8) (9). This adaptability enables it to exploit a wide array of resources, ensuring its nutritional needs are met even in nutrient-poor environments (10–12). Its short generation time and ability to be reared in large numbers under controlled laboratory conditions allows precise manipulation of diet, environmental factors, and genetic backgrounds, leading to reproducible results and the ability to conduct high-throughput screening. Studying this species thus provides a unique window into fundamental mechanisms of nutrient digestion and absorption, ecological flexibility, and the optimization of nutrient cycling within biological systems. By meticulously controlling dietary macronutrient profiles—specifically the protein-to-carbohydrate (P:C) ratio—we can simultaneously investigate the black soldier fly’s life history traits, host gene expression, and the associated microbial community’s structure and gene expression. This comprehensive approach sheds light on the precise mechanisms underlying this organism’s remarkable resilience and offer broad implications for understanding how organisms, including humans, thrive within diverse nutritional landscapes.

## Results

### Black Soldier Fly Life-History Traits: Larval Weight, Survivorship and Days to Pupation

Overall, diet composition significantly affected black soldier fly larval weight (one-way ANOVA: F = 20.43; df = 2; *p* = <0.001) (Figure 1A), survivorship (one-way ANOVA: F = 5.603; df = 2; *p* = 0.010) (Figure 1B), and days to pupation (one-way ANOVA: F = 3457; df = 2; *p* = <0.001) (Figure 1C). Pairwise comparisons (Tukey’s HSD) determined significant differences in final larval weight between treatments. When reared on the equal ratio diet, larval weight was 28.3% greater (*p* < 0.01) then when raised on the carbohydrate-biased diet, and 19.7% greater than larvae fed the protein-biased diet (*p* < 0.01, Figure 1A). Additionally, larval survivorship was negatively impacted (*p* = 0.025) on the protein-biased diet, with a 22.9% decrease in survivorship compared to the carbohydrate-biased diet, and a 23.8% decrease (*p* = 0.018) in survivorship compared to the equal ratio diet (Figure 1B). Furthermore, significant differences in timing to black soldier fly pupation between treatments was observed. The number of days for larvae to reach the 40% pupation threshold was negatively impacted (*p* ≥ 0.01) on the carbohydrate-biased diet (Figure 1C). Larvae required 75.8% longer time to pupate compared to the equal ratio diet and 77.2% longer time to pupate compared to the protein-biased diet (*p* < 0.01) (Figure 1). In summary, the equal ratio diet (1P:1C) resulted in the optimal performance among the three diets utilized in this study, as larvae exhibited the highest weight and survivorship with no reduced time to pupation (Figure 1).

**Figure 1.**
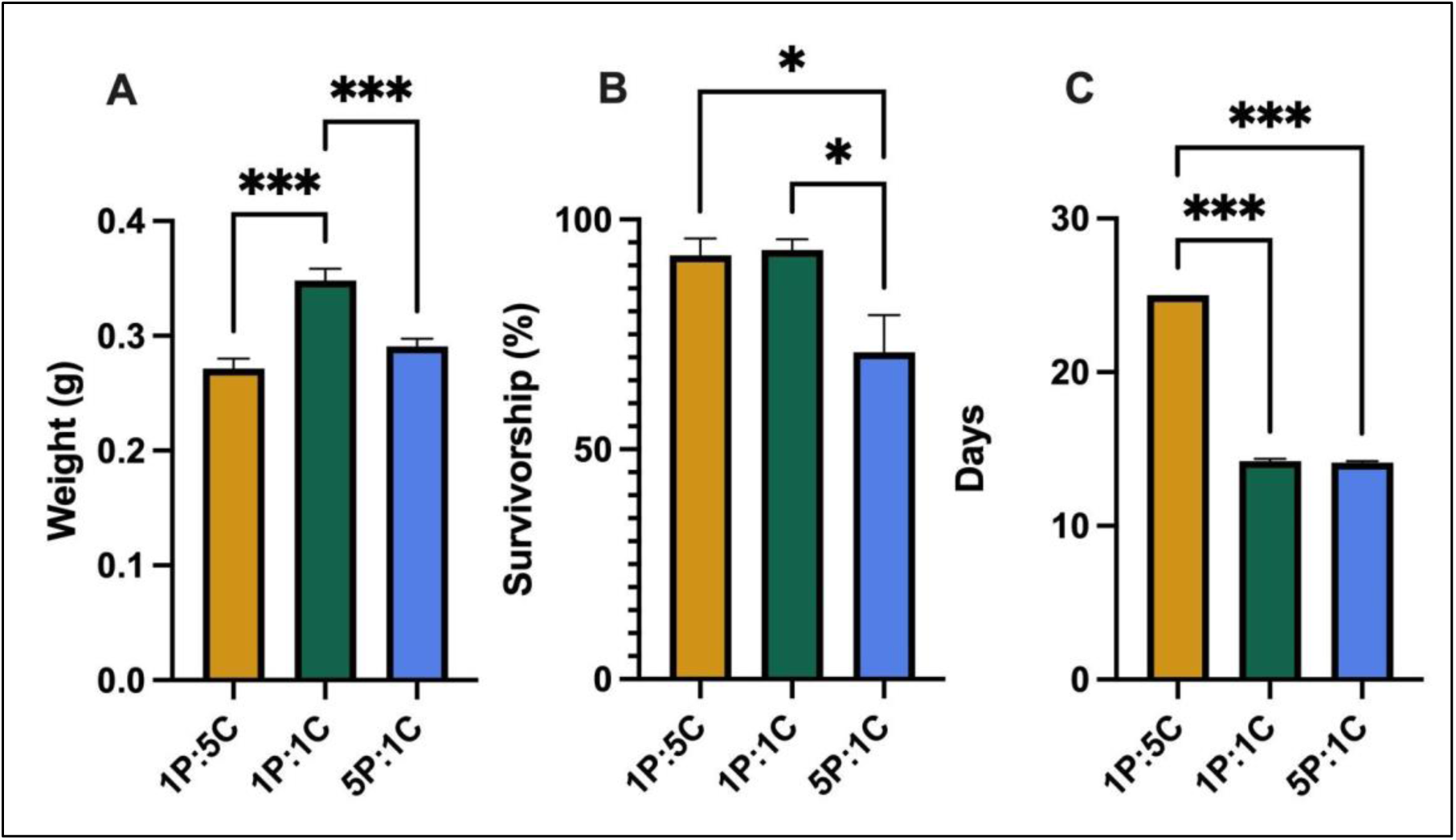
Measurements of larval life history traits. A) Average larval weight (g) +/- SEM; B) survivorship +/- SEM, and; C) days to 40% prepupation +/- SEM for larvae reared on either a carbohydrate biased (1P:5C), balanced (1P:1C), or protein biased (5P:1C), artificial diet. Significance following a Tukey’s HSD post-hoc test was determined (* *p* = 0.0, ** *p* = 0.00, *** *p* ≤ 0.001).

### Black Soldier Fly Gene Expression

Following rRNA removal, an average of 55,729,300 reads were generated per sample (range: 9,993,402 - 151,186,941). Of these, an average of 54,429,300 reads (range: 97.9% −98.5%) were successfully mapped to the black soldier fly reference genome. The reads corresponded to a total of 15,574 transcripts, of which 13,492 met the read count criteria outlined in the methods and were included in subsequent analyses. A total of 401 significantly differentially expressed transcripts across the three diet treatments were identified using a likelihood ratio test (LRT) with a Benjamini-Hochberg adjusted *p*-value cutoff of 0.05. These differentially expressed genes (DEGs) were subsequently categorized into four groups (A-D) based on diet-dependent expression patterns (Figure 2).

**Figure 2.**
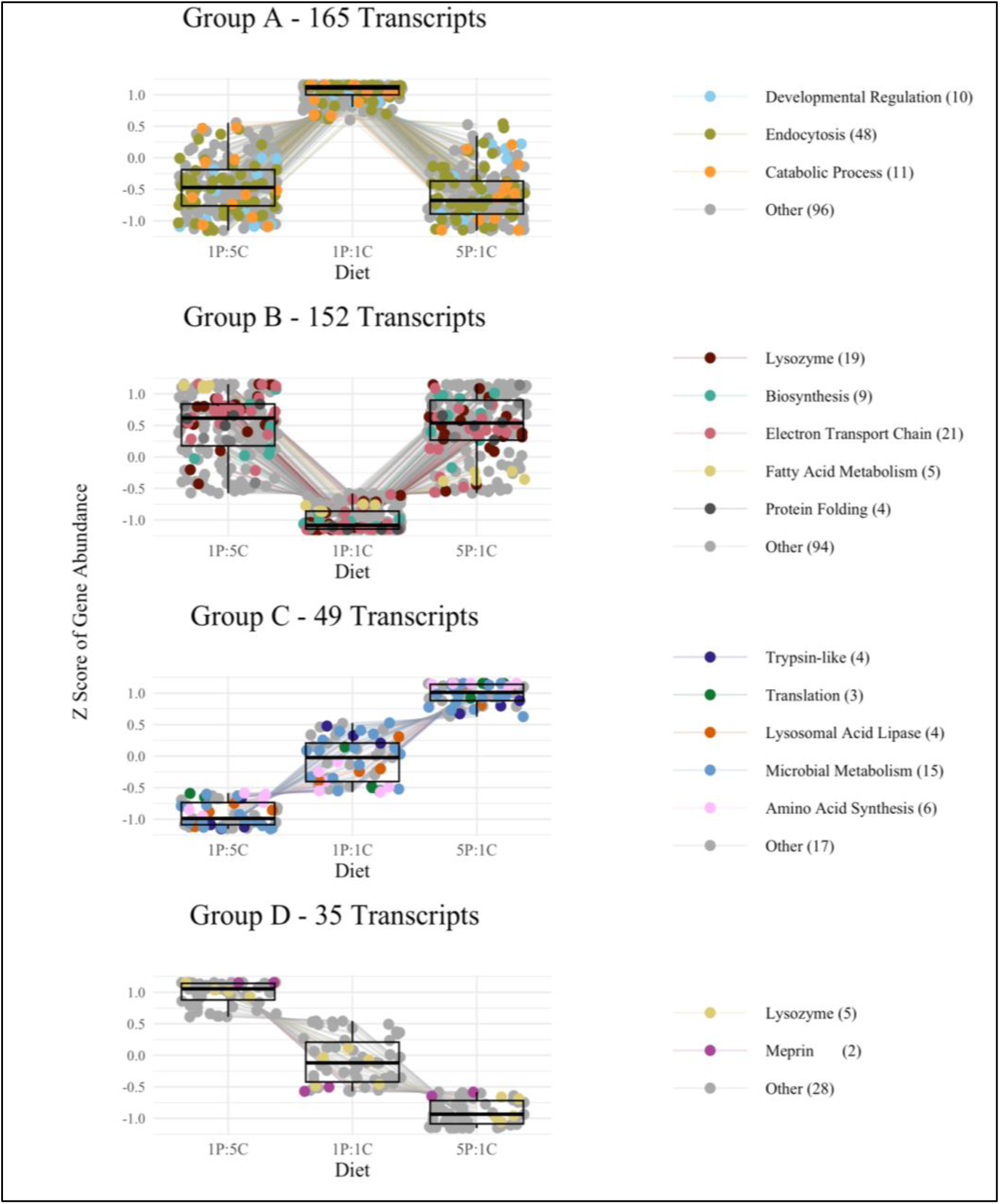
Clustering of transcripts from BSF larval gut tissue based on expression profile across diet treatments and a summation of functional enrichment analysis. Differentially expressed transcripts across three diet treatments were clustered into four groups (A-D) based on expression profiles using DEGreport by means of divisive analysis clustering. Expression profiles for each group are depicted (left) where z-score of gene abundance is shown on the y-axis and diet treatment is shown on the x-axis. Based on enriched KEGG pathway and ontology terms as well as GO biological function terms, summary terms were manually curated to avoid redundancy and better represent the gene sets and depicted for each group (right). See Data S1 for a complete list of functional enrichment results.

Groups A and B comprised 165 and 152 transcripts, respectively, accounting for 79% of all significantly differentially expressed transcripts. In Group A, transcripts were upregulated in the balanced diet relative to the other diets, while Group B transcripts exhibited the opposite pattern, being downregulated in the balanced diet compared to the other diets. Conversely, Groups C and D, consisting of 49 and 35 transcripts, respectively, contained transcripts whose expression demonstrated a more linear relationship with the P:C ratios of the diet treatments. Specifically, in Group C, significant expression increased as the diets progressed from carbohydrate-biased, to balanced, to protein-biased, whereas Group D exhibited the opposite trend.

To elucidate the enriched functions and pathways within each expression group, significant terms were identified utilizing the Gene Ontology (GO) (biological process) and Kyoto Encyclopedia of Genes and Genomes (KEGG) (pathway and ontology) databases. Enriched terms were subsequently manually curated to provide a more comprehensive summary of each transcript and reduce redundancy within and between terms. Data S1 contains the complete functional enrichment results for each group, as well as the manually curated terms displayed in Figure 2. Several terms enriched in Group A encompassed Catabolic Processes (10 transcripts), Developmental Regulation (48 transcripts), and Endocytosis (11 transcripts) (Figure 2A). Enriched Group B transcripts comprised those associated with Biosynthesis (19 transcripts), Electron Transport Chain (9 transcripts), Fatty Acid Metabolism (21 transcripts), Lysozyme (4 transcripts), Mitochondrial Transport (5 transcripts), and Protein Folding (4 transcripts) (Figure 2B). Group C was characterized by enrichment in Amino Acid Transport, Lysosomal Acid Lipase, Microbial Metabolism, Translation, and Trypsin-like activities, consisting of four, three, four, fifteen, and six transcripts, respectively (Figure 2C). Finally, Group D comprised six transcripts coding for lysozyme proteins and two transcripts coding for metalloproteinase (meprin) proteins (Figure 2D).

### Microbial Community Analysis

Overall, 2,985 taxa were assigned across all samples from the Kraken2/Bracken pipeline and used for community analysis. Diet composition significantly affected alpha diversity in both the larvae gut microbiome (one-way ANOVA: *p* = 8.4e-05, Additional File 1: Table S1) and the frass microbiome (one-way ANOVA: *p* = 4.6e-04, Additional File 1: Table S2). Using pairwise t-tests, we found that alpha diversity was significantly different between the carbohydrate-biased diet and both the balanced and protein-biased diet treatments (p-adjusted = 0.001 for both comparisons) in the larval microbiomes (Figure 3A, Additional File 1: Table S3). However, between the balanced ratio and protein-biased diet, alpha diversity was no different (*p* = 0.791) (Figure 3A, Additional File 1: Table S3). Interestingly, opposite patterns were observed in the frass microbiome diversity with the carbohydrate-biased diet having significantly lower diversity, than the balanced ratio (*p* = 3.66e^-4^) and the protein-biased diets (*p* = 0.013) (Figure 3B, Additional File 1: Table S4). However, alpha diversity between the balanced ratio diet and the protein-biased diet were not different in the frass (*p* = 0.532, Additional File 1: Table S4).

**Figure 3:**
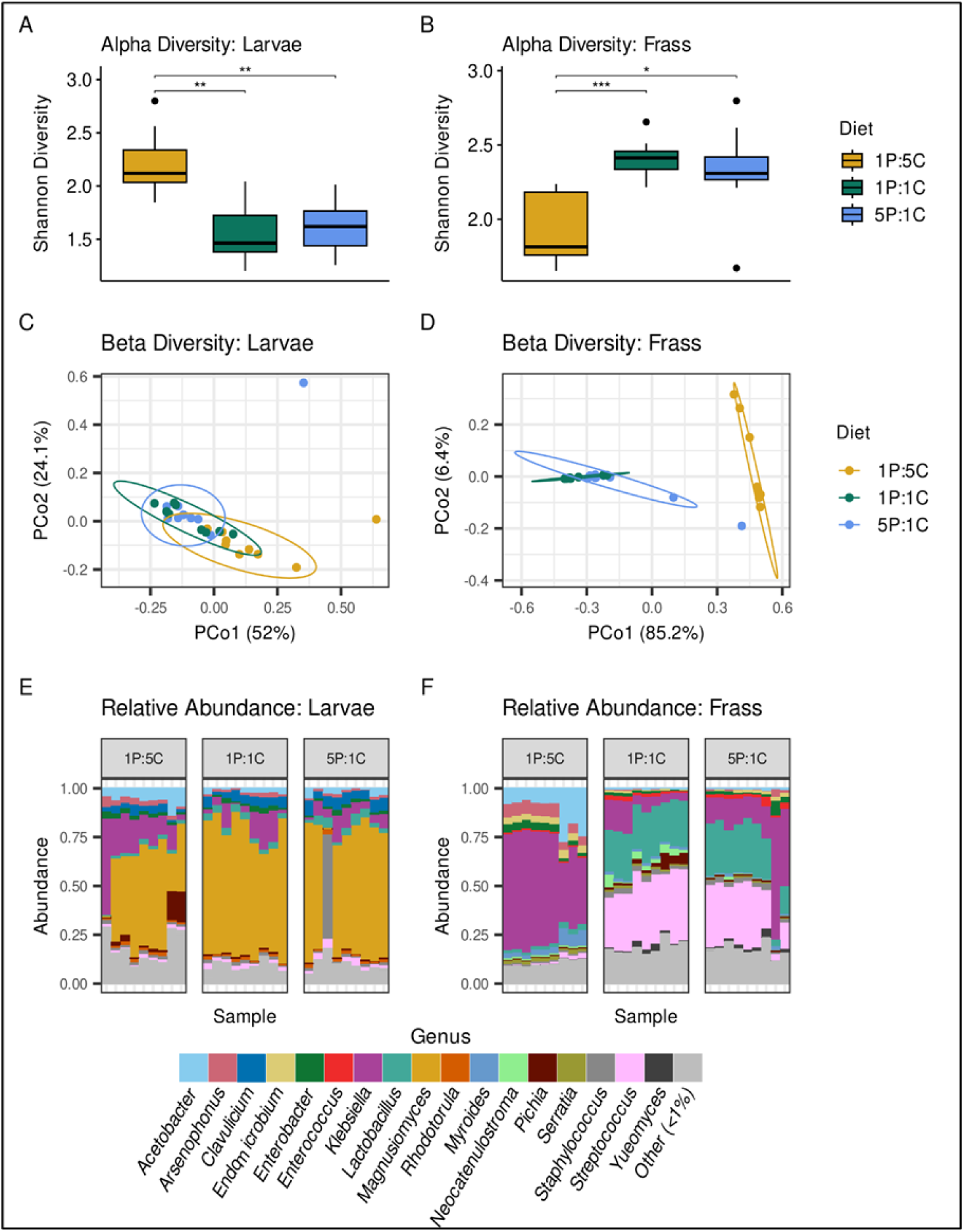
Diversity of microbial communities within the BSF larvae (A&C) and frass (B&D). **(A-B)** Alpha diversity between different diet groups was measured using the Shannon Diversity index (SDI). Comparisons of SDI between diet compositions were conducted using pairwise T-tests. Only statistically significant comparisons are labeled. **(C-D)** Beta diversity was measured using Bray-Curtis dissimilarities. PERMANOVA analysis showed significant differences in overall microbial taxonomic composition between diets within larvae (p = 0.00014, 10^5^ permutations) and frass (p < 1e-05, 10^5^ permutations). **(E-F)** Relative microbial abundance for larvae and frass, respectively. Taxa that had an overall average less than 1% relative abundance for larvae and frass are not shown.

Principal coordinate analysis (PCoA) was employed to evaluate the variation in community composition among individual samples (Figure 3C, 3D). The effect of diet composition was more pronounced in the frass microbiome compared to the larvae microbiome, with the first principal coordinate accounting for 85.2 % of the variance in the frass and distinguishing the carbohydrate-biased diet from the other two diets (Figure 3D). In the larvae, the impact of diet composition on the microbiome was less substantial, as evidenced by continuous variation and overlapping clusters. However, 76.1 % of the variance was explained by the first two principal coordinates (Figure 3C). To determine whether overall community composition differed significantly between diet treatments, we first confirmed that beta diversity group dispersions were homogeneous within the treatments (Permutation Test: *p* = 0.525 and *p* = 0.647 for larvae and frass, respectively, Tables S5 and S6). Although differences in larval microbial diversity between samples were not apparent in the PCoA (Figure 3C), significant differences in overall community composition based on diet composition were observed in both larvae and frass samples (PERMANOVA: *p* = 1.4e-04 and *p* = 1e-05 for larvae and frass microbiomes, respectively, Tables S7 and S8). Pairwise comparisons revealed that differences in overall microbial community in the larvae were attributable to the carbohydrate-biased diet versus the equal ratio and protein-biased diets (*p* = 0.003 and *p* = 0.006, respectively), while the equal ratio and protein-biased diets did not differ significantly from each other (*p* = 1) (Additional File 1: Table S9). However, in the frass, all diets had significantly different microbial composition from each other (Additional File 1: Table S9).

When we investigated the relative abundance of microbial taxa, we found that most of the larval gut microbiome was made up of fungal taxa specifically from the *Magnusiomyces* genus (Figure 3E). On the other hand, dominant taxa in frass microbiomes were primarily bacterial. In the carbohydrate-biased diet, *Klebsiella* was generally the most dominant taxa, while *Streptococcus*, *Lactobacillus*, and *Klebsiella* comprised most of the microbial communities within the equal ratio and the protein-biased diets (Figure 3F). Although the equal ratio and protein-biased frass microbiomes formed overlapping clusters in our PCoA analysis (Figure 3D), pairwise comparisons showed significant differences between the microbial community of all three diet treatments (Additional File 1: Table S9).

### Microbial Metatranscriptomic Analysis

The initial metatranscriptome assemblies resulted in a total of 1,061,655 and 1,597,113 microbial transcripts for the larvae and frass metatranscriptomes, respectively. After clustering and filtering, the number of transcripts in our assemblies was reduced to 10,273 transcripts in the larval assembly, and 132,729 transcripts in the frass assembly. We compared the expression levels of these transcripts across each diet treatment and found 498 differentially expressed transcripts overall. These differentially expressed transcripts were clustered into groups based on their expression patterns across treatments (Figure 4, Data S2). For the larval transcripts, this resulted in three groups. Group A consisted of 439 transcripts (∼88%) that were upregulated in the carbohydrate-biased diet but downregulated in the equal ratio and the protein-biased diets (Figure 4A). Group B consisted of 22 transcripts (∼4%) that were downregulated in the equal ratio diet, upregulated in the protein-biased diet, but were not dysregulated in the carbohydrate-biased diet (Figure 4B). Group C consisted of 37 transcripts (∼7%) that were downregulated in the carbohydrate-biased diet compared to the equal ratio and protein-biased diets (Figure 4C).

**Figure 4:**
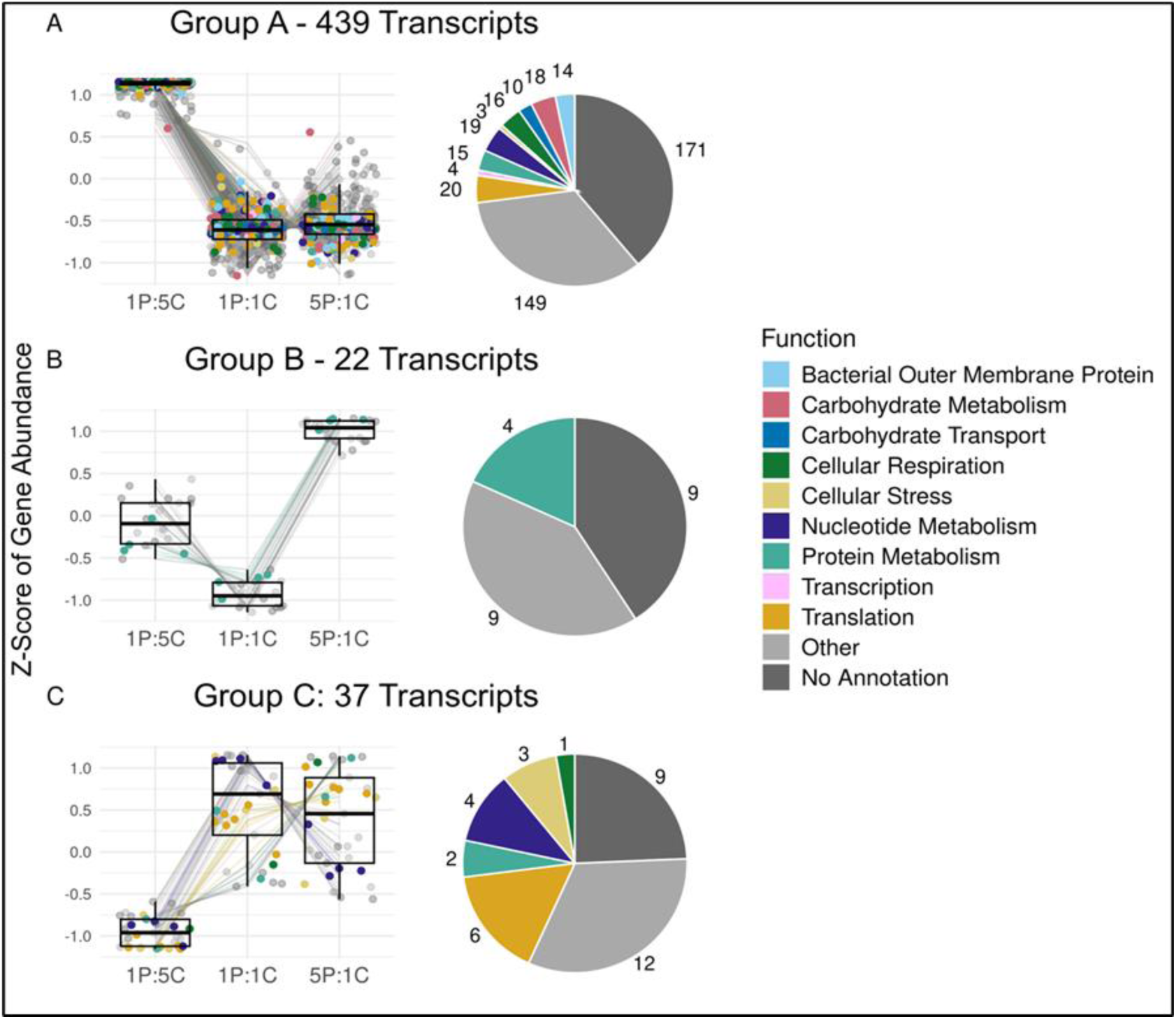
Functional enrichment of differentially expressed genes in the BSF larvae gut microbiome. (A-C) Differentially expressed transcripts were grouped by expression patterns across diet treatments. Functional profiles were created using functional annotations from eggNOG, and significantly enriched functional terms from Gene Ontology, KEGG Orthology, and KEGG Pathway (Data S2).

Using functional annotations and enrichment analyses, we found that many differentially expressed genes matched expected functions within their diet composition. In group A, (upregulated in carbohydrate-biased, downregulated in other diets) we found 18 microbial genes involved in carbohydrate metabolism and transport (Figure 4A). Additionally, other cellular processes were enriched in this group such as components of cellular respiration and nucleotide metabolism (Figure 4A). Although there were fewer transcripts overall in groups B and C, we found 4 annotated transcripts associated with protein metabolism upregulated in the protein-biased diet. Overall, many microbial transcripts had uninformative functional annotations or were difficult to categorize (Figure 4). Functional assessment was particularly challenging in the frass, as there were 20,433 differentially expressed transcripts. These differentially expressed transcripts formed four groups based on expression patterns across diet treatments with hundreds to thousands of enriched GO terms within these groups. These tables are provided in Data S3.

### Microbial and Black Soldier Fly Correlations

To determine if there were changes in black soldier fly gene expression associated with diet-induced changes of the microbiota, we correlated to larval microbiome alpha diversity metrics (Shannon Diversity Index) with normalized gene expression counts of the differentially expressed black soldier fly genes. We found eight of the 399 differentially expressed black soldier fly genes significantly correlated (Spearman’s Rho, Bonferroni *p*-adjusted value < 0.05) to larval microbiome alpha diversity (Table 2). Most of these black soldier fly genes are predicted to be involved in digestion and immunity (13–17). Additionally, we determined whether relative abundance of microbial taxa was correlated to abundance of differentially expressed black soldier fly genes at each taxonomic rank, but we did not find any significant correlations.

To investigate whether specific microbial genes were associated with expression patterns in black soldier flies, we also correlated the abundance of differentially expressed microbial transcripts with the differentially expressed black soldier fly genes. This resulted in 12 microbial transcripts (one was removed due to high sequence similarity to a black soldier fly gene) with significant correlations to 10 black soldier fly genes (Figure 5). Interestingly, four of the 10 black soldier fly genes that were correlated with microbial transcripts were also correlated to larval microbiome alpha diversity, including two involved in pathogen recognition and multiple that are involved in digestion (Table 2, Figure 5) (13–17).

**Figure 5:**
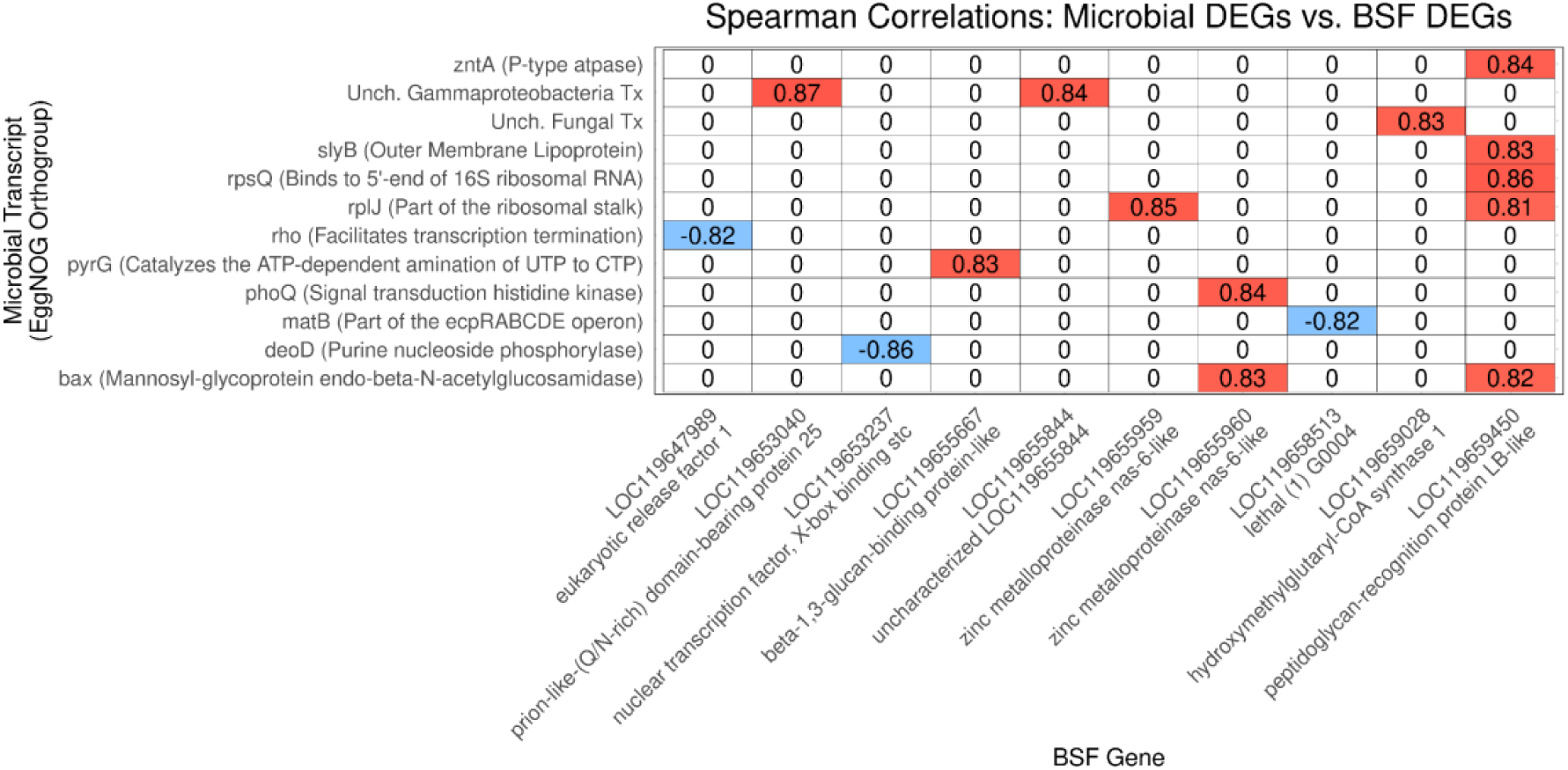
Microbial transcript expression influences BSF gene expression. Only genes with significant (Bonferroni p-adjusted < 0.05) correlations are shown, and non-significant correlations were changed to zeros. Microbial transcripts are labeled with their “preferred names” and “description” from EggNOG mapper. BSF genes are labeled with their NCBI geneID, along with their description.

## Discussion

The intricate relationship between what an organism eats, the microbes residing within its gut, and its overall health is a fundamental biological principle with profound implications across the animal kingdom, including for humans. Our study, focusing on the highly adaptable black soldier fly, offers valuable insights into how dietary composition orchestrates changes in the gut microbiome and, in turn, influences host fitness. These findings resonate with broader ecological and physiological concepts, highlighting the delicate balance required for optimal health.

In this study, RNA sequencing was employed to measure host-diet-microbe tripartite interactions of a generalist species, the black soldier fly. In addition to measuring black soldier fly life-history traits, RNA sequencing was utilized because (1) the transcriptome serves as a robust proxy for an organism’s physiological state, and (2) it enables simultaneous sequencing of the transcriptome of black soldier fly and its associated microbial community. This approach facilitated the exploration of the dynamic interplay between the host and its microbiome by investigating how diet composition influences host fitness and gene expression, while concurrently measuring how the associated microbial community changes and functionally responds to these same conditions.

The black soldier fly serves as a model organism to provide insights into how a highly resilient, generalist feeder adapts to and utilizes macronutrients in response to dietary fluctuations. Furthermore, this research is of interest to black soldier fly scientists and producers, as both black soldier fly larvae and their associated microbes contribute to the insect’s unique bioconversion capabilities, which is also why numerous sectors of the food and feed industry are interested in mass rearing them (18). Overall, we observed that manipulating diet P:C ratios significantly affected black soldier fly life history traits, gene expression, and the black soldier fly-associated microbial community and transcript expression. Although other studies have assessed changes in the black soldier fly transcriptome or metatranscriptome caused by diet (10, 19, 20), this is the first study to measure changes in both host gene expression and microbiome metatranscriptomic changes in response to carefully targeted changes in macronutrient profile within the same sample.

Black soldier fly larvae fed the balanced diet exhibited the most substantial number and most pronounced patterns of differential gene expression (Figure 2A, 2B). Among the black soldier fly life-history traits measured, the balanced ratio diet demonstrated optimal performance, resulting in the highest larval weight, highest survival rate, and shortest time to pupation (Figure 1).

Notably, this outcome was reflected in black soldier fly gene expression, as functional categories associated with developmental regulation were enriched in the set of genes upregulated in the balanced diet treatment (Figure 2A). A balanced diet provides the most diverse array of nutrients that may serve as cofactors and signaling molecules, directly influencing the expression of genes involved in various cellular processes. Consequently, these genes may be responsible for the increased weight of the larvae fed the balanced diet. Moreover, in the set of genes upregulated in the unbalanced diets, biosynthesis-related genes were enriched, suggesting that black soldier fly larvae expend energy to supplement their dietary requirements in the carbohydrate- and protein-biased diets, which could account for their reduced weight. Although differences in black soldier fly gene expression and fitness were observed between the balanced diet and the nutrient-biased diets, the microbiomes were most similar between the balanced diet and the protein-biased diet, indicating that microbes were not the sole factor contributing to the black soldier fly’s success on this diet.

Although both the carbohydrate-biased (1P:5C) and protein-biased (5P:1C) diets exerted adverse effects on the life history of the black soldier fly, examining larvae and microbial responses to these diets can elucidate how black soldier flies and other hosts adapt to suboptimal nutritional conditions (10). When subjected to a carbohydrate-biased diet, larvae exhibited reduced weight and prolonged development time, while larval survival remained unaffected. This carbohydrate-biased diet likely induced protein deficiency, impairing tissue development and consequently hindering larval ability to construct essential structures for pupation. Additionally, the carbohydrate-biased diet may have influenced energy allocation and protein utilization. Notably, the carbohydrate-biased diet had the most pronounced impact on the microbial community associated with the black soldier fly: (1) microbial diversity in the larval gut was significantly elevated compared to other diets (Figure 3A); and (2) the largest group of differentially expressed transcripts in the larval gut microbes were upregulated under the carbohydrate-biased diet (Figure 4A). This heightened activity and diversity may be attributed to carbohydrates serving as a superior energy source and providing a wider array of substrates for microbes in the black soldier fly gut than other macronutrients (21). Furthermore, the functions represented in the differentially expressed microbial transcripts suggest that the microbes actively participate in metabolizing the carbohydrate-rich diet. For instance, carbohydrate metabolism and carbohydrate transport are enriched in the upregulated microbial transcripts in the carbohydrate-biased diet, along with cellular respiration, the process by which organisms derive energy from the breakdown of carbohydrates (Figure 4A).

Nevertheless, the observed increase in microbial diversity and function within the carbohydrate-biased diet may be linked to the decreased performance of black soldier fly larvae on this diet, potentially due to nutrient piracy or the metabolic costs associated with immunity (22, 23). Supporting this hypothesis, significant positive correlations were identified between alpha diversity and the expression of genes encoding pathogen-recognition receptors (Table 2).

Moreover, the expression of these pathogen recognition genes was also correlated with the expression of certain microbial transcripts (Figure 5). This indicates that as microbial populations expand within the gut, black soldier flies may enhance their surveillance mechanisms and nutritional immunity against potential microbial pathogens, possibly expending additional energy to manage opportunistic microbes. Notably, another study reported that black soldier fly larval samples exhibiting higher alpha diversity also demonstrated reduced larval weight gain (24).

Additionally, we observed that a set of lysozyme genes was upregulated in the black soldier fly gut when reared on the carbohydrate-based diet compared to the other diets (Figure 2B, 2D). Lysozymes play a role in immunity as antimicrobial effectors that degrade peptidoglycan in bacterial cell membranes. Interestingly, it has been noted that detritivores, which naturally consume a microbe-rich diet, can utilize lysozyme enzymes to digest bacteria in their gut, ultimately absorbing the nutrients released post-digestion (25–27). This alternative nutrient acquisition route may explain why black soldier fly larvae reared on the carbohydrate-biased diet did not exhibit decreased survival.

Conversely, a negative correlation was observed between the microbial gene *matB* and black soldier fly gene *lethal* (1) G0004. The *matB* gene encodes a structural protein of the *Escherichia coli* pilus, facilitating adherence to the intestinal mucosa (28). Upregulation of *matB* may enhance *E. coli* colonization in the insect gut, potentially leading to dysbiosis that could impair host digestion, nutrient absorption, and immune function, thereby negatively regulating “lethal” genes essential for black soldier fly fitness and survival. Another example of complex interactions between host-microbiome-diet is the observed upregulation of the microbial *rho* concomitant with the downregulation of the *eukaryotic release factor 1* (*erf1*) (Figure 5). The *rho* gene is involved in bacterial gene regulation, controlling the expression of microbial genes, while *erf1* is crucial for translation termination. These data indicate black soldier fly larvae may limit the synthesis of certain proteins or metabolites that its commensal microbes depend upon, creating a competitive environment where the host regulates access to essential resources. This suggests that hosts, such as black soldier flies, may influence the metabolic functions of their microbial partners by strategically supplying or restricting specific nutrients or growth substrates (29, 30). Although this proposed mechanism is intriguing, it remains speculative until further experimental data can provide support. While the concept of nutritional provisioning is not novel, the precise mechanisms by which a host regulates specific nutrient availability to target particular microbial species or metabolic pathways, the molecular mechanisms involved in fine-tuning this control, and the potential evolutionary trade-offs remain open and exciting areas for future research.

The protein-biased (5P:1C) diet adversely affected the black soldier fly life-history resulting in decreased survival rates and larval weight (Figure 1). Analysis of black soldier fly gene expression revealed that genes upregulated exclusively in the protein-biased diet were associated with protein metabolism (Figure 2C). This includes genes responsible for protein catabolism, such as trypsin-like genes, and genes involved in protein anabolism, including amino acid synthesis genes and translation genes (Figure 2C). Similarly, upregulated transcripts related to protein metabolism were observed in the microbial community, albeit to a lesser extent (Figures 4B, 4C). These findings suggest that both black soldier fly larvae and microbes are metabolizing the abundant dietary proteins. However, the microbial community composition and gene expression profiles were comparable between the balanced ratio diet and the protein-biased diet (Figures 3 and 4). Consequently, it remains unclear why the protein-biased diet resulted in significant reductions in larval weight and survival. One plausible explanation is that the high-protein diet may have led to increased levels of ammonia or other toxic byproducts within the insect and substrate environment, thereby reducing larval fitness and survival, although these factors were not investigated in this study.

Another notable finding from this study was the substantial fungal abundance observed in the larvae microbiome data. While numerous studies have concentrated on bacterial components in black soldier fly microbiome research, there is a relative paucity of investigations into the mycobiome. Within the black soldier fly gut, the fungal genus *Magnusiomyces* emerged as the most dominant taxon across all treatments (Figure 3). Additionally, three other fungal genera—*Pichia*, *Clavulicium*, and *Rhodotorula*—were among the top taxa present in the larval gut.

Previous studies examining the fungal community of black soldier flies have generally identified *Pichia* and *Candida* species as the predominant fungal taxa (31–33). Although *Magnusiomyces* has not been previously reported in the black soldier fly microbial community, its family, Dipodascaceae, has been documented in black soldier fly gut tissues (33). A prior study reported a high abundance of an unidentified *Geotrichum* species in larval guts, which also belongs to the Dipodascaceae family, and some *Magnusiomyces* species were formerly classified as *Geotrichum* species (34). Notably, species of both *Magnusiomyces* and *Pichia* are known to engage in mutualistic relationships with larvae of other dipteran insects, detoxifying harmful compounds from decaying plant matter for cactophilic *Drosophila* (35). Given that wild black soldier fly larvae typically consume decomposing material, these fungal taxa may perform a similar function in black soldier flies. Furthermore, positive correlations were found with microbial *pyrG*, which plays a crucial role in pyrimidine biosynthesis, and a black soldier fly gene identified as *beta 1,3 glucan binding protein-like,* potentially involved in recognizing fungal organisms. The *pyrG* gene in fungi is well-studied and frequently utilized as a genetic marker (36). In addition to fungal taxa in the larval guts, a high abundance of bacterial genera such as *Klebsiella* and *Acetobacter* was observed, which have been identified as dominant bacterial genera associated with black soldier flies in previous studies (37, 38). Importantly, the results suggest that nutrient utilization in the black soldier fly gut is a multi-kingdom effort, as 141 of the 498 differentially expressed microbial transcripts were assigned to eukaryotic orthogroups, with 69 of these further classified as fungal (31) (Figure S1).

The frass samples revealed a distinct impact of diet composition on the microbiome, particularly in the carbohydrate-biased (1P:5C) diet (Figure 3B, 3D, 3F). Notably, *Klebsiella* emerged as the predominant genus in the carbohydrate-biased diet, whereas *Lactobacillus* and *Streptococcus* species were more prevalent in the balanced (1P:1C) and protein-biased (5P:1C) diets (Figure 3E). This observation aligns with previous studies on black soldier fly substrates (39–41). In contrast to the larval gut microbiome, fungal taxa did not dominate the frass microbial community. However, in the 1P:5C control diet (1P:5C diet run alongside the experiment but without larvae), the fungal genus *Mucor* was the dominant microbial taxon (Figure S2). Interestingly, *Mucor* was no longer dominant when larvae were present (Figure 3F), corroborating a previous study indicating that black soldier fly larvae reduce the fungal diversity of their substrate (42).

Furthermore, this suggests that the high representation of *Magnusiomyces* in the larval gut is not due to its high presence in the diet. Nevertheless, *Magnusiomyces* was detected at low relative abundance in diet samples without larvae (<1% average across all control diets), indicating that even if present at low levels in black soldier fly substrate, this fungus can proliferate in the larval gut.

Overall, we observed three key patterns in the black soldier fly that carry significant implications for our understanding of host-microbe-diet interactions. First, a carbohydrate-rich diet led to a notable reduction in black soldier fly weight and a shorter pupation period, while simultaneously boosting microbial diversity and activity within the gut. This intriguing effect may suggest a biological trade-off, where an energy-rich, yet perhaps nutritionally imbalanced, diet could lead to altered resource allocation between host metabolism and immune function. The observed correlation between increased microbial diversity and specific insect immune genes hints at a dynamic interplay, where the host might be investing resources in managing a more diverse microbial community, potentially at the expense of growth (43). This echoes observations in human health, where certain dietary patterns can influence gut microbial diversity and immune responses, sometimes leading to inflammatory states (44).

Second, a protein-rich diet had a detrimental impact on black soldier fly larval survival and weight, yet surprisingly, it did not significantly alter the gut microbiome composition relative to the balanced diet. This suggests that the negative effects were likely due to either insufficient overall nutrition or the accumulation of toxic metabolic byproducts from excessive protein, rather than direct microbial dysbiosis. The lack of significant microbial shifts between the protein-rich and balanced diets is particularly noteworthy, suggesting that the host’s physiological limits, rather than dramatic microbial community changes, were the primary drivers of decreased performance in this specific dietary context. This finding underscores the importance of considering host-intrinsic factors alongside microbial dynamics when assessing dietary impacts, a crucial consideration in human nutrition where excessive intake of certain macronutrients can lead to metabolic stress independent of immediate microbiome alterations (45).

Third, the balanced diet consistently yielded the most favorable outcomes across all assessed life-history traits. This optimal performance is likely attributable to its provision of a comprehensive range of nutrients, thereby minimizing the energetic demands on the host for metabolic regulation and potentially reducing the need for extensive microbiome management. This highlights a universal principle: a well-balanced diet provides the necessary substrates for both host and microbe to thrive synergistically. While our study in black soldier flies offers broad insights into the resilience of a generalist detritivore to macronutrient variations, the precise molecular mechanisms underlying this resilience warrant further investigation through functional studies. However, these findings provide a valuable framework for understanding the intricate interplay between diet, microbes, and host health, offering potential avenues for future research into human dietary interventions and the optimization of gut health.

Understanding the intricate interplay between host genetics, microbial communities, and dietary influences is paramount for developing personalized health strategies and unraveling the complex etiology of numerous diseases; therefore, continued research into this tripartite relationship is essential for advancing both basic science and clinical applications. The black soldier fly is a promising model to unravel the nuanced interactions within the host-microbe-diet triad and could be a crucial study system for optimizing health, enhancing resilience to environmental stressors, and ultimately, ensuring the long-term well-being and evolutionary success of animal populations, including humans.

## Materials and Methods

### Black Soldier Flies

Black soldier fly larvae were obtained from EVO Conversion Systems, LLC, College Station, Texas, USA and transferred to the Forensic Laboratory for the Investigative Entomological Sciences (F.L.I.E.S.) at Texas A&M University, College Station, Texas, USA for the duration of the experiment. The larvae were 7-day-old neonates that had been fed Gainesville diet (consisting of 50% wheat bran, 20% corn meal, and 30% ground alfalfa, (46)) at 70% moisture.

### Diet Formulation

Three dietary treatments were formulated in a nutrition laboratory at Texas A&M University (Table 1). The treatment formulations varied in their protein-to-carbohydrate ratios and were designated as follows: 1P:5C (carbohydrate-biased), 1P:1C (balanced protein:carbohydrate), and 5P:1C (protein-biased). The relative moisture content ratios and mean weights of the dry diet formulations utilized in the experiments, as varying quantities of water were required to achieve a consistent feed texture across diet formulations (Additional File 1: Table S10). Diets were prepared according to methods described in previous insect-nutrition-based studies (47–49).

**Table 1:** Microbial diversity is correlated with some differentially expressed genes in BSF. DE group refers to the group each gene was assigned to in. **Figure 2**.

| Gene | Spearman's Rho | Bonferroni $p$-adjusted | NCBI Description | Putative Function | DE Group |
| --- | --- | --- | --- | --- | --- |
| LOC119655959 | 0.80 | $p < 0.001$ | zinc metalloproteinase nas-6-like | Protease | D |
| LOC119659450 | 0.74 | $p = 0.004$ | peptidoglycan-recognition protein LB-like | Immunity (pathogen recognition) | D |
| LOC119655667 | 0.73 | $p = 0.006$ | beta-1,3-glucan-binding protein-like | Immunity (pathogen recognition) | B |
| LOC119661453 | -0.74 | $p = 0.007$ | nascent polypeptide associated complex protein alpha subunit | Protein homeostasis | C |
| LOC119657638 | -0.73 | $p = 0.012$ | RNA exonuclease prage | RNA degradation | A |
| LOC119651252 | 0.71 | $p = 0.014$ | testicular acid phosphatase homolog | Digestion | B |
| LOC119655960 | 0.70 | $p = 0.017$ | zinc metalloproteinase nas-6-like | Protease | D |
| LOC119654621 | 0.69 | $p = 0.029$ | zinc metalloproteinase nas-6-like | Protease | B |

**Table 2:**
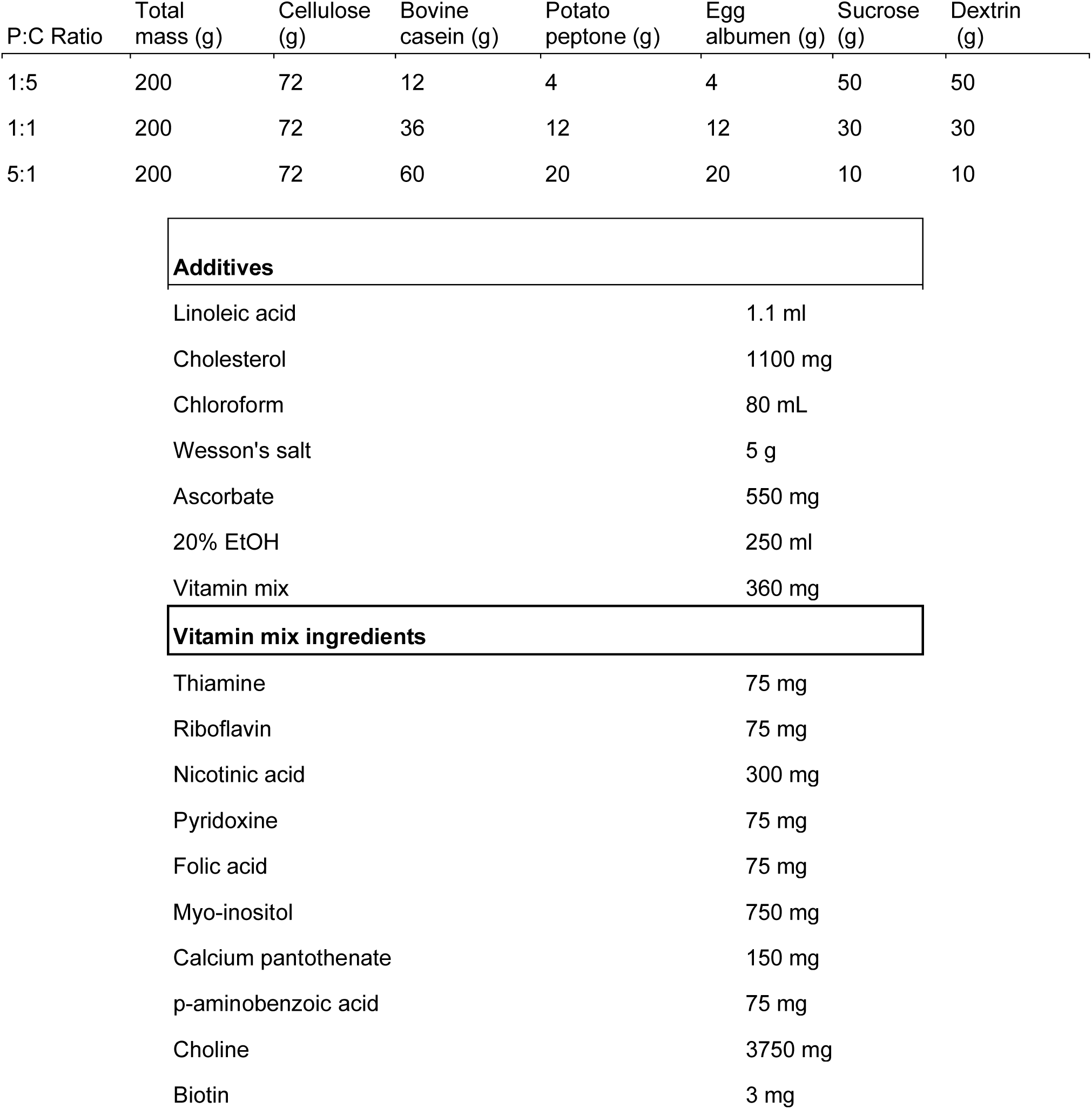
Artificial diet formulations at 60% macronutrient content with respective adjusted P:C ratios. Protein ingredients (i.e., bovine casein, potato peptone, and egg albumen) are consistently present in a 3:1:1 ratio, while carbohydrate ingredients (i.e., sucrose and dextrin) are consistently present in a 1:1 ratio. The additives enumerated below remain constant across all dietary treatments irrespective of the P:C ratio.

| P:C Ratio | Total mass (g) | Cellulose (g) | Bovine casein (g) | Potato peptone (g) | Egg albumen (g) | Sucrose (g) | Dextrin (g) |
| --- | --- | --- | --- | --- | --- | --- | --- |
| 1:5 | 200 | 72 | 12 | 4 | 4 | 50 | 50 |
| 1:1 | 200 | 72 | 36 | 12 | 12 | 30 | 30 |
| 5:1 | 200 | 72 | 60 | 20 | 20 | 10 | 10 |

Additives
|  |  |
| --- | --- |
| Linoleic acid | 1.1 ml |
| Cholesterol | 1100 mg |
| Chloroform | 80 mL |
| Wesson's salt | 5 g |
| Ascorbate | 550 mg |
| 20% EtOH | 250 ml |
| Vitamin mix | 360 mg |

Vitamin mix ingredients
|  |  |
| --- | --- |
| Thiamine | 75 mg |
| Riboflavin | 75 mg |
| Nicotinic acid | 300 mg |
| Pyridoxine | 75 mg |
| Folic acid | 75 mg |
| Myo-inositol | 750 mg |
| Calcium pantothenate | 150 mg |
| p-aminobenzoic acid | 75 mg |
| Choline | 3750 mg |
| Biotin | 3 mg |

### Feeding Experiment

For each diet treatment and control, twenty 50 ml Falcon® centrifuge tubes (Corning®, Glendale, AZ, USA) were allocated with 40 ml of their respective diets. Ten tubes from each diet treatment were inoculated with 10, 7-day-old black soldier fly larvae. A paper towel was placed on top of each tube for ventilation and secured with a rubber band to prevent larval escape. The remaining 10 tubes for each diet treatment were left uninoculated for subsequent analysis of variation in microbial communities resulting from diet treatment tubes with versus without larvae present. Experimental tubes were maintained in an incubator (Percival, Geneva Scientific, Fontana, WI, USA) at 25 °C, 70% humidity, and on a 12:12 light-dark cycle. All tubes inoculated with larvae were inspected daily for indications of pre-pupation (e.g., darkening of the cuticle). Entire tubes with four or more larvae beginning to darken were removed and subsequently harvested.

Corresponding diet tubes not inoculated with larvae were also harvested at that time to maintain consistency for microbial analysis. The resulting larvae were weighed individually using an Adventurer Pro™ balance (OHAUS CORP., NJ, USA). Each larva was placed in individual VWR® Microcentrifuge Tubes (Avantor®, PA, USA) and labeled accordingly, while the remaining diet was returned to its originally labeled tube for microbial analysis. All samples were then stored in a freezer at −20 °C.

### Black Soldier Fly Larvae Dissections and Residual Sampling

Methods for sampling, RNA Isolation, library preparation, sequencing, and metatranscriptome assembly are described in (50). Three falcon tubes were randomly selected for each diet composition, and three black soldier fly larvae were haphazardly sampled from each tube. Larval cuticles were sterilized by briefly immersing whole larvae in 70% ethanol followed by a rinse with sterile water. Sterilized larvae were immediately placed in RNAlater (Invitrogen, Waltham, MA, USA) and whole gut tissues were dissected using sterilized forceps (70% ethanol spray) under a Zeiss SteREO Discovery microscope (Zeiss, Oberkochen, Germany). Individual whole larval guts were placed in 1.5 mL of RNAzol (Molecular Research Center, Cincinnati, OH, USA) for RNA isolation. Additionally, Black soldier fly residual (frass) was sampled by homogenizing the contents of individual falcon tubes with a sterile laboratory spatula and collecting 1 gram of frass, which was immediately placed in 1.5 mL of RNAzol. This procedure was repeated three times for each tube.

### Total RNA Isolation

Samples were homogenized in RNAzol utilizing a Genolyte 1200 tissue homogenizer (Spex SamplePrep, Metuchen, NJ, USA) at 4,000 RPM with 3 mm autoclave-sterilized stainless-steel beads. The homogenate was centrifuged for 5 minutes at 12,000 x g, and 1 mL of the supernatant was extracted for subsequent RNA purification. The manufacturer’s protocol for RNA isolation using RNAzol was followed through the precipitation of DNA, polysaccharides, and protein steps. Subsequently, the aqueous portion containing RNA was transferred directly into an NEB Monarch RNA cleanup kit (New England Biolabs, Ipswich, MA, USA). To complete RNA isolation, the NEB’s “Purification of RNA from the Aqueous Phase Following TRIzol/Chloroform Extraction using the Monarch RNA Cleanup Kits” protocol was implemented. RNA purity was assessed utilizing a NanoDrop One (Thermo Fisher Scientific, Grand Island, NY, USA) spectrophotometer, and concentration was determined using a Qubit 2.0 fluorometer (Invitrogen, Waltham, MA, USA). RNA integrity was evaluated using a 4150 TapeStation System (Agilent, Santa Clara, CA, USA).

### Library Preparation and Sequencing

Whole shotgun metatranscriptome sequencing libraries were prepared using the NEB Ultra II RNA kit, with the omission of rRNA depletion and mRNA enrichment steps to preserve all host and microbial reads. The resultant cDNA was multiplexed using NEBNext Oligos for Illumina (New England Biolabs, Ipswich, MA, USA), thereby generating unique barcodes for each library. The quality of the libraries was evaluated using a 4150 TapeStation System, and their concentration was quantified using a Qubit 2.0 fluorometer. Subsequently, the uniquely barcoded libraries were pooled according to diet. The pooled libraries were sequenced by Psomagen on a HiSeq 2000 instrument (Illumina, San Diego, CA, USA), generating paired 151 bp reads.

### Quantification and Statistical Analyses

#### Insect Gene Expression Analyses

Adapter sequences and low-quality bases were trimmed from the reads using Trim Galore 4.5 (51) prior to assessing read quality using FastQC 11.9 using default parameters (52). Ribosomal RNA (rRNA) reads were removed by aligning the reads with reference rRNA sequences (accessions: XR_005250514, XR_005250512, XR_005249317, XM_038046990, XM_038068750) utilizing the Burrows–Wheeler transform algorithm with default settings (BWA 7.12, (53)), retaining the unmapped reads. The unmapped filtered reads were subsequently aligned to the black soldier fly reference genome (GCF_905115235.1, (54)) using BWA default settings, followed by sorting and indexing with Samtools 1.15 (55). Transcript counts were then quantified using HTSeq 2.0.3 (56).

Differential gene expression analysis was conducted in R 4.3.0 (57) utilizing DEseq2 1.40.2 (58). To maintain data quality, transcripts with total read counts less than 10 were excluded from the analysis. A likelihood ratio test (LRT) was performed to identify transcripts whose expression varied significantly across the three diet treatments. Genes were considered differentially expressed if they met the Benjamini-Hochberg (BH) adjusted *p*-value threshold of 0.05 and a base mean expression of five. The differentially expressed transcripts were subsequently clustered into groups based on expression profiles across the three diet treatments using DEGreport 3.18 (59) and the divisive analysis clustering algorithm (60). Lists of transcripts found in each group were then utilized for functional enrichment analysis.

The reference black soldier fly proteome was annotated with Kyoto Encyclopedia of Genes and Genomes (KEGG) and Gene Ontology (GO) terms utilizing the web-based eggnog-mapper (61). Protein sequences were annotated based on orthology to sequences in the EggNOG 5 database employing e-score, bit-score, percent identity, and query coverage thresholds of 0.001, 60, 40 %, and 20%, respectively. KEGG pathway and ontology enrichment was conducted using R with clusterProfiler 4.8.2 (62); significant terms were generated using a hypergeometric test and a BH adjusted *p*-value threshold of 0.05. In contrast, GO – biological function enrichment was conducted using the topGO 2.52.0 (63) package, and significant terms were identified using Fisher’s exact test and the weight01 algorithm with a *p*-value threshold of 0.05. Based on the results from these functional enrichment analyses, summary terms were manually curated to minimize redundancy and more accurately represent the transcripts.

### Read QC, Filtering, and Mapping for Microbial Analyses

Read quality was assessed using FastQC v0.11.9 (52). Low-quality bases and adapters were removed from the reads using Trimmomatic v.0.39 in paired-end mode using the ILLUMINACLIP option to remove Illumina universal adapters, a sliding window of 4:15, and a minimum length of 30 (64). Paired, trimmed reads were then mapped to the black soldier fly reference genome (GCF_905115235.1, (54)) using HISAT2 v.2.2.1(65), directing unmapped reads to new fastQ files using the “un-conc” option. Reads that did not map to the black soldier fly genome were used for microbial analyses.

### Microbial Community Analysis

To assign microbial taxonomy, non-black soldier fly reads were mapped to release 138 of the SILVA small subunit rRNA database (66) utilizing Kraken2 v2.1.2 in paired-end mode (67). Taxonomic abundance was estimated using Bracken v2.8 at every taxonomic level up to class level with a threshold of 10 reads required to estimate abundance (68). Each sample’s Bracken report was compiled into one BIOM-format file using kraken-biom at each taxonomic level (69), and the genus level-abundance estimates were utilized for further analysis. This file was converted to a phyloseq object in R using the import_biom command from the phyloseq (v1.42.0) package (57, 70). Alpha diversity was measured using Shannon’s Diversity Index obtained from the estimate_richness command in phyloseq. Differences in alpha diversity between diet compositions were assessed using a one-way ANOVA and pairwise T-tests with Holm-Bonferroni corrections for multiple comparisons. Beta diversity was measured using Bray-Curtis dissimilarity metrics and visualized using principal coordinate analysis (PCoA) from the plot_ordination command in phyloseq. Relative abundance within each sample was calculated using the abundance estimates from Bracken divided by the sum of the abundance of all taxa. Group dispersions within diet treatments were measured by providing the Bray-Curtis dissimilarity matrix to the betadisper function in the VEGAN R package (71). Homogeneity of dispersions within the treatment groups was evaluated using the permutest function using 999 permutations. Statistical differences in overall community composition were assessed using Bray-Curtis dissimilarity and permutational multivariate ANOVA (PERMANOVA) with 100,000 permutations. Pairwise comparisons of treatments following PERMANOVA were conducted using the pairwise.perm.manova command from the RVAideMemoire v0.9-83-7 package in R (72) employing the Bonferroni method to account for multiple comparisons.

### Microbial Metatranscriptome Assembly and Annotation

Microbial metatranscriptomes were assembled for each sample utilizing Trinity v.2.14.0 via their docker container (73, 74). Metatranscriptome assemblies were subsequently clustered by larval or frass samples employing CD-HIT-EST with a 90% similarity threshold, a word count of 8, and a minimum length of 500 nt (75). Subsequently, the representative transcript from each cluster underwent functional annotation using eggNOG mapper v.2.1.9 in DIAMOND mode, employing the “metagenome” input type and --allow_overlaps set to “none” (61). Following this process, all transcripts annotated as “Metazoa” or “Viridiplantae” were filtered out. Only the annotation with the lowest e-value was retained when transcripts possessed multiple annotations.

### Microbial Gene Expression Analyses

Transcript abundance was estimated by pseudo-mapping the non-black soldier fly reads to the representative transcripts after metatranscriptome clustering and filtering using Kallisto v0.44.0 (76). Abundance estimates were imported to R using tximport v1.26.1 (77), and differential expression analysis was conducted using DEseq2 v1.38.3 employing the likelihood ratio test (58). Patterns of gene expression across treatment groups were detected using the degPatterns command in the DEGreport package v1.34.0 (59). Only genes with an adjusted *p*-value (BH) less than 0.05 and a mean base expression level greater than 5.0 were considered. Gene Ontology (GO) enrichment analyses were conducted using topGO v2.50.0 (63), and Kyoto Encyclopedia of Genes and Genomes (KEGG) orthology and pathway enrichment analyses were conducted using clusterProfiler v4.6.2 (62) with annotations assigned by eggNOG. Genes assigned to groups by the degPatterns command were further manually annotated with a “summary term” based on their functional annotation, biological process GO term enrichment, KEGG orthology enrichment, and KEGG pathway enrichment.

### Correlations

Correlations were conducted utilizing matrices constructed from normalized black soldier fly gene expression counts (from DESeq2) and alpha diversity estimates (see previous sections) for each sample. Only differentially expressed genes were considered. Spearman’s rho between matrices was determined using the associate function within the R package “microbiome” v 1.28.0 in “table” mode with the method set to “spearman,” the p.adjust_method set to “bonferroni,” p.adj.threshold set to 0.05, and the minimum number of significant correlations set to one.

Correlations between the normalized counts of black soldier fly differentially expressed genes and the normalized counts of differentially expressed microbial transcripts were conducted using the same parameters. Furthermore, we correlated relative abundance of all taxa at each taxonomic rank and the normalized counts of differentially expressed black soldier fly genes using the same criteria, but no significant results were returned.

## Acknowledgments

The authors would like to acknowledge the Center for Insect Biomanufacturing and Innovation for supporting this work.

## Funding

The work presented here was supported by funding from industry advisory board members of the Center for Insect Biomanufacturing and Innovation, and by support from: National Science Foundation grant 2052454 National Science Foundation grant 2052565 National Science Foundation grant 2052788 Any opinions, findings, and conclusions or recommendations expressed in this material are those of the author(s) and do not necessarily reflect the views of the National Science Foundation.

## Supporting Information

**SI Appendix**

Figures S1–S2 and Table S1-S10

**Dataset Files**

Dataset 1: Excel file containing additional data too large to fit in a PDF, related to Figure 2.

Dataset 2: Excel file containing additional data too large to fit in a PDF, related to Figure 4.

Dataset 3: Excel file containing additional data too large to fit in a PDF, frass associated transcripts.

## Notes

### Competing Interest Statement

The authors have declared no competing interest.

