## Supplemental Tables and Figures for "A Perfect Soldier: the black soldier fly as a microbial-mediated physiological resilience model"

**This PDF file includes:**

Figures S1 to S2

Tables S1 to S10

**Other supporting materials for this manuscript include the following:**

Data S1 to S3

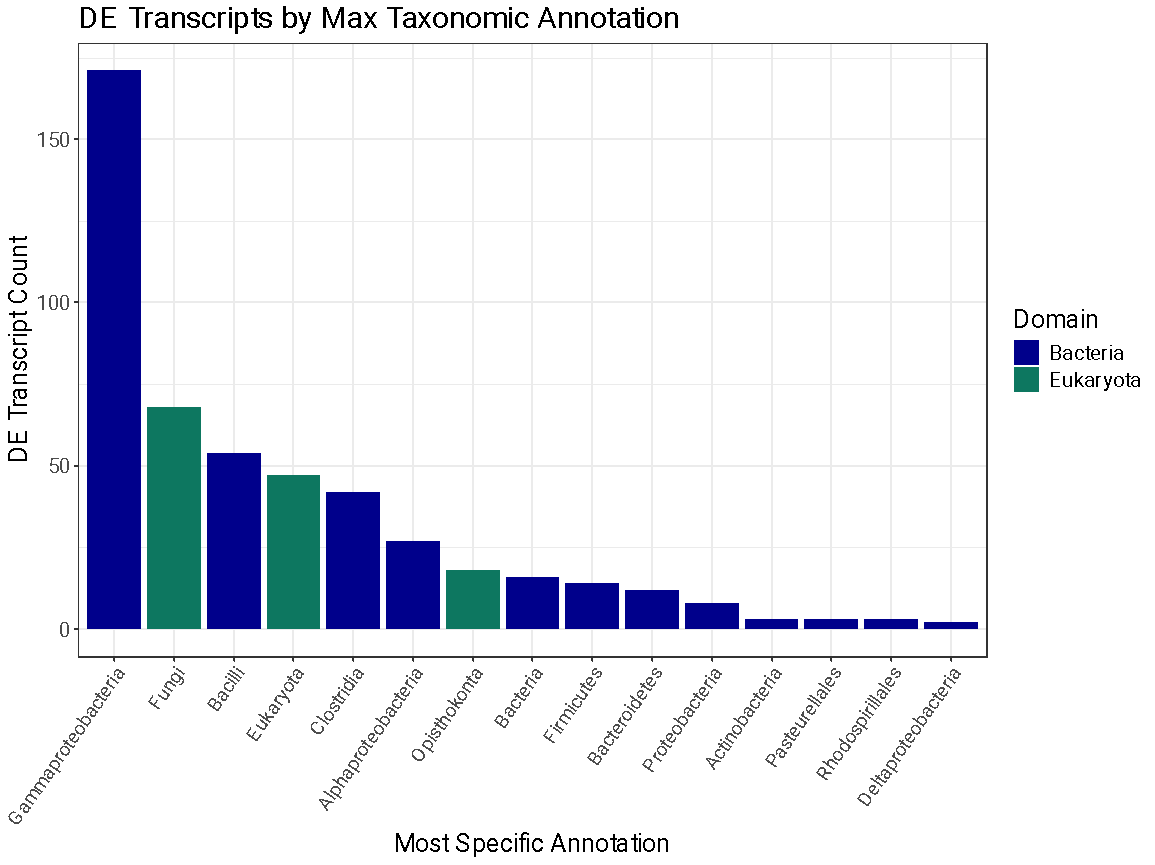

**Fig. S1.**

Taxonomic representation of differentially expressed microbial transcripts.

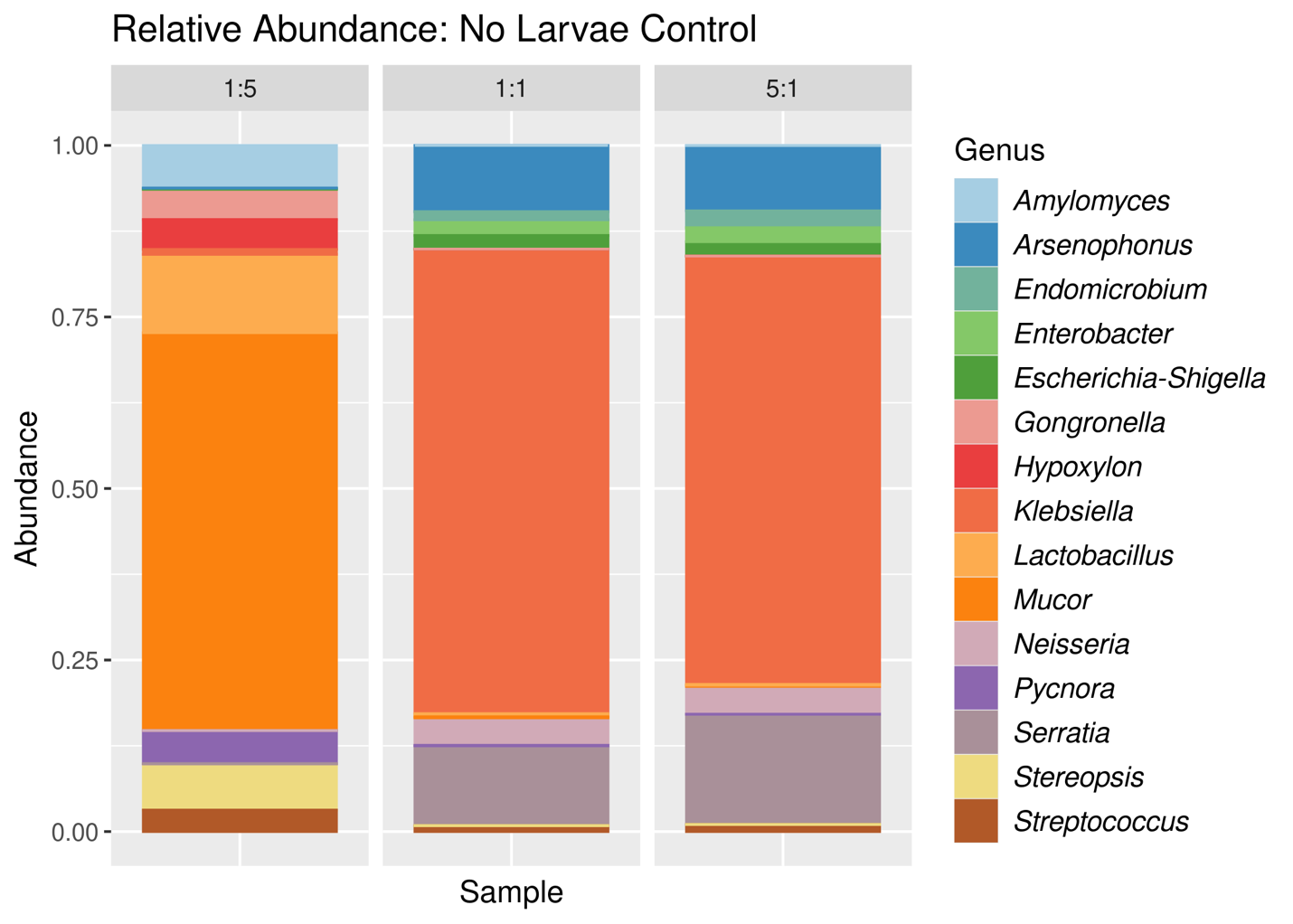
**Fig. S2.**

Control diet microbial community. Top 15 taxa present in the control diets. Other taxa had a relative abundance of <1%.

**Table S1: ANOVA table for larval gut alpha diversity (Shannon diversity index).**

|  | Df | Sum of Sqs | Mean sq | F | p |
| --- | --- | --- | --- | --- | --- |
| Diet | 2 | 2.314 | 1.1572 | 14.23 | 8.4e-05 |
| Residuals | 24 | 1.952 | 0.0813 |  |  |

**Table S2: ANOVA table for frass alpha diversity (Shannon diversity index).**

|  | Df | Sum of Sqs | Mean sq | F | p |
| --- | --- | --- | --- | --- | --- |
| Diet | 2 | 1.194 | 0.5971 | 10.75 | 0.000464 |
| Residuals | 24 | 1.333 | 0.0556 |  |  |

**Table S3: Pairwise T-tests of larval microbiome alpha diversity (Shannon diversity index).** Adjustments for multiple comparisons were conducted using the Holm-Bonferrni method. Significant comparisons are in bold.

| Diet 1 | Diet 2 | n1 | n2 | statistic | Df | p | p.adj |
| --- | --- | --- | --- | --- | --- | --- | --- |
| 1:5 | 1:1 | 9 | 9 | 4.51 | 16.0 | 3.59e-4 | 0.001 |
| 1:5 | 5:1 | 9 | 9 | 4.55 | 15.4 | 3.61e-4 | 0.001 |
| 1:1 | 5:1 | 9 | 9 | -0.269 | 15.7 | 0.791 | 0.791 |

**Table S4: Pairwise T-tests of frass alpha diversity (Shannon diversity index).** Adjustments for multiple comparisons were conducted using the Holm-Bonferrni method. Significant comparisons are in bold.

| Diet 1 | Diet 2 | n1 | n2 | statistic | Df | p | p.adj |
| --- | --- | --- | --- | --- | --- | --- | --- |
| 1:5 | 1:1 | 9 | 9 | -5.40 | 13.0 | 1.22e-4 | 3.66e-4 |
| 1:5 | 5:1 | 9 | 9 | -3.16 | 14.7 | 7e-3 | 1.3e-2 |
| 1:1 | 5:1 | 9 | 9 | 0.645 | 10.9 | 0.532 | 0.532 |

**Table S5: Permutation test for homogeneity of diet group dispersions in the larval gut microbiome.** Results were determined using Bray-Curtis dissimilarity and 999 permutations.

|  | Df | Sum of Sqs | Mean sq. | F | p |
| --- | --- | --- | --- | --- | --- |
| Diet Group Distance | 2 | 0.03945 | 0.019724 | 0.704 | 0.514 |
| Residuals | 24 | 0.67243 | 0.028018 |  |  |

**Table S6: Permutation test for homogeneity of diet group dispersions in the frass microbiome.** Results were determined using Bray-Curtis dissimilarity and 999 permutations.

|  | Df | Sum of Sqs | Mean sq. | F | p |
| --- | --- | --- | --- | --- | --- |
| Diet Group Distance | 2 | 0.02176 | 0.010879 | 0.4483 | 0.665 |
| Residuals | 24 | 0.58236 | 0.024265 |  |  |

**Table S7: PERMANOAVA results for larval gut microbiome community.** Results were determined using Bray-Curtis dissimilarities and 100,000 permutations.

|  | Df | Sum of Sqs | R2 | F | p |
| --- | --- | --- | --- | --- | --- |
| Diet | 2 | 0.53253 | 0.27454 | 4.5413 | 0.00014 |
| Residuals | 24 | 1.40719 | 0.72546 |  |  |

**Table S8: PERMANOAVA results for frass microbiome community.** Results were determined using Bray-Curtis dissimilarities and 100,000 permutations.

|  | Df | Sum of Sqs | R2 | F | p |
| --- | --- | --- | --- | --- | --- |
| Diet | 2 | 3.1972 | 0.75678 | 37.338 | 1e-05 |
| Residuals | 24 | 1.0275 | 0.24322 |  |  |

**Table S9: Pairwise PERMANOAVA results for larvae and frass following PERMANOVA.** Results were determined using Bray-Curtis dissimilarities with 999 permutations. Corrections for multiple testing were performed using the Bonferroni method.

|  | 1P:5C vs 1P:1C | 1P:5C vs 5P:1C | 1P:1C vs 5P:1C |
| --- | --- | --- | --- |
| Larvae | 0.003 | 0.009 | 1 |
| Frass | 0.003 | 0.003 | 0.009 |

**Table S10: Moisture content ratios and mean weights of dietary mixtures utilized in the experimental procedures.** p10:c50: low-protein diet; p30:c30: balanced diet; p50:c10: high-protein diet.

| **Diet** | **Moisture %** | **Avg dry weight (g)** |
| --- | --- | --- |
| p10:c50 | 45.32% dry, 54.68% wet | 42.3 g |
| p30:c30 | 39.47% dry, 60.53% wet | 37.1 g |
| p50:c10 | 43.28% dry, 65.72% wet | 34.5 g |

**Data S1. (separate file)**

Excel file containing additional data too large to fit in a PDF, related to Figure 2.

**Data S2.** **(Separate file)**

Excel file containing additional data too large to fit in a PDF, related to Figure 4.

**Data S3. (Separate file)**

Excel file containing additional data too large to fit in a PDF, frass associated transcripts.
